# Assessment of CD4 T Lymphocyte Cell Levels among Hepatitis B, C and E Viruses Negative Individuals in Ibadan, southwestern Nigeria

**DOI:** 10.1101/185165

**Authors:** M. O. Adewumi, E. C. Omoruyi, I. M. Ifeorah, A. S. Bakarey, A. O. Ogunwale, A. Akere, T. O. C. Faleye, J. A. Adeniji

**Affiliations:** Department of Virology, College of Medicine, University of Ibadan, Ibadan, Nigeria.; Institute of Child Health, College of Medicine, University of Ibadan, Ibadan, Nigeria.; Department of Medical Laboratory Sciences, College of Medicine, University of Nigeria, Nsukka, Nigeria.; Institute for Advanced Medical Research & Training, College of Medicine, University of Ibadan, Ibadan, Nigeria.; Oyo State College of Agriculture and Technology, Igboora. Oyo State, Nigeria.; Department of Medicine, College of Medicine, University of Ibadan, Ibadan, Nigeria.; Department of Microbiology, Faculty of Science, Ekiti State University, Ado Ekiti. Nigeria.

**Author notes:** Corresponding author Adewumi O.M, Omoruyi E.C., Ifeorah I.M., Bakarey A.S., Ogunwale A.O., Akere A., *Faleye T.O.C., Adeniji J.A.

**Keywords:** CD4 T lymphocyte, HBV, HCV, HEV, HIV, Nigeria

## Abstract

The CD4 T lymphocytes play a key role in achieving a regulated effective immune response to foreign antigens. It is also a valuable parameter for assessing HIV disease progression. However, variations in CD4 T lymphocyte values due to diverse factors have been reported. We evaluated CD4 T lymphocytes among healthy community dwellers who tested negative for hepatitis B, hepatitis C and hepatitis E viruses and compared the results with the National Reference Values (NRVs). Four hundred consenting participants who fulfilled the criteria for enrolment were evaluated for CD4 T lymphocyte counts. Estimated mean CD4 T lymphocyte count of 1,183 (CD4 Range: 328-2680) cells/μl of blood was recorded for the participants. Four (1.0%), 151 (37.8%), 157 (39.2%), 74 (18.5), and 14 (3.5) of the participants had CD4 T lymphocyte count ranged 352-500, 501-1,000, 1,001-1500, 1501-2,000, and >2,000 cells/μl of blood, respectively. Differences in the estimated mean CD4 count between different age groups varied significantly (P=0.010). In this study, significantly higher CD4 T lymphocyte values were observed among the study population in comparison to the NRVs, and consequently we advise careful interpretation and use of extrapolated CD4 T lymphocyte values in the management of persons with diverse geographical background or health conditions.

## Introduction

The immune system is solely responsible for the recognition and subsequent elimination of foreign antigens, in addition to other vital functions [1]. Specifically, the lymphocyte cells, mainly made up of the T-lymphocytes (thymus-derived), B-lymphocytes (bone-marrow-derived), and the NK (natural killer) cellsfunction cooperatively to achieve effective immune response[1,2]. NaiveCD4 T lymphocyte cells are activated and differentiated into distinct effector subtypes depending mainly on the cytokine milieu of the microenvironment, and the newly differentiated cells subsequently play major role in mediating immune response through the secretion of specific cytokines[2]. The cells carry out multiple functions, ranging from activation of B cells for antibody production, recruitment and activation of macrophages, and recruitment of neutrophils, eosinophils, and basophils to sites of infection and inflammation[1]. Precisely, the CD4 T lymphocytes play a key role in achieving a regulated effective immune response to foreign antigens.

Regrettably, some viruses specifically target and inhibit the CD4 T lymphocyte cellfunctions[3]. Notably, the Human Immunodeficiency Virus (HIV) infects CD4 T lymphocyte cells selectively, and causesits destruction directly, as well as indirectly, thus, leading to gradual loss of the cells in the peripheral circulation[4].Consequently, monitoring of the CD4 T lymphocyte cell count became an important parameter for assessing HIV disease progression, and it hassince remained probably the most important marker of immune dysfunction in HIV infection. It isfrequently used to decide the initiation of antiretroviral therapy (ART), to monitor the efficacy of ART and to start treatment for opportunistic infections (OIs), especially in resource-limited settings.

Reference CD4 T lymphocyte values were established in developed countries and extrapolated for use in clinical settings in other regions of the world. Such reference values help in proper assessment of the degree of immune deficiency during HIV disease progression. Over time, variations were observed in CD4 T lymphocyte values among different populations of the world [5-7], and several factors[5, 8-13] were identified. Specifically, infection with some viral pathogens including HIV [14] and hepatitis viruses [2, 15-17] havebeennoted to influencecharacteristics ofCD4 T lymphocyte cells.

However, despite the reported specifics and documented variations in CD4 T lymphocyte cell levels, consequent of the factors mentioned,data from developed countries werelong used to determine the threshold levels of CD4 T lymphocyte counts in developing countries. In Nigeria, attempts were made by researchers in different regions of the country to determineacceptable reference values for CD4 T lymphocytes count among Nigerians. Ultimately, a country-wide study was conducted by Oladepo et al. [18] to determine CD4 T lymphocyte cell levels among Nigerians, and this was subsequently referred to as the National Reference Values (NRVs) in this manuscript. Enrollees for the country-wide CD4 evaluation study were defined as apparently healthy persons. The ‘apparently healthy’ participants were so defined based on their HIV negative status alone, without consideration for their HBV, HCV or HEV status in a region ofknown hepatitis endemicity. Of note, infection with any of these often result in chronic viral hepatitis, and such individuals could have passed as ‘healthy’ at enrolment without precise screening for hepatitis viruses. Further, emergence of exhausted T lymphocytes with reduced effector properties has been reported during chronic hepatitis infections[3, 19, 20]. Thus,the possible contributions of the hepatitis viruses in T lymphocyte quality and functions cannot be underestimated.

Further, bearing in mind that Nigeria is a very heterogeneous country, variation in CD4 T lymphocyte values is expected, and that emphasizes the need for the establishment of reference CD4 T lymphocyte values by regions of the country. Consequently, this study was designed to determine, compare and evaluate the reference values of CD4 T lymphocytes among healthy residents of Ibadan who tested negative for hepatitis B, hepatitis C and hepatitis E viruses.

## Methodology

### Study Population

This study was a part of a larger viral hepatitis study carried out by the Evolutionary Dynamics of Hepatitis in Nigeria (EDHIN) group in 2013. Out of a total of 1,115 participants enrolled for the larger study, only 437 participants who consented to participate in the CD4 evaluation study were included in this study.

### Study Location

A cross-sectional study was conducted among healthy residents of Ibadanmetropolis. Ibadan city,the capital of Oyo State in the southwestern region of Nigeria is one of the most populous cities in the country, witha population of over 3 million people. It covers a total area of 3,080 square kilometers, making it the country’s largest city by geographical area. Ibadan metropolis enjoys optimal distribution of primary, secondary and tertiary health institutions. Specifically, Ibadan is home to the University College Hospital, the foremost tertiary institution in Nigeria and other secondary healthcare facilities.

### Enrolment of Participants

The study participants were enrolledusing a convenient sampling technique between July and September, 2013. Specifically, attendees at the 2013 World Hepatitis Day lecture at the University College Hospital, Ibadan were counselled on the need for regular medical examinations and subsequently offered free medical tests.Semi-structured questionnaire which captured socio-demographic characteristics was administered on each of the 437 consenting participants. Thereafter, blood samples were collected from each participant for laboratory investigations. As a fall out of the lecture, interested members of the community visited the Institute of Child Health, College of Medicine, University College Hospital for free medical examination. During their visits, prospective participants were also counselled on the benefits of regular medical examinations, andindividually administered questionnaire to capture demographic information. Thereafter, blood sample was collected from each consenting participant for laboratory investigations.

### Sample Collection, Analysis and Storage

Five milliliters of blood specimen was collected from each participant by venipuncture. The blood sample was immediately dispensed into an appropriately labeled tube containing ethylene diamine tetra acetic acid (EDTA), and subsequently transported to the laboratory in the Institute of Child Health, College of Medicine, University of Ibadan, in a sample box with ice packs to maintain cold temperature. For participants who visited the institute, blood samples were collected and immediately submitted for laboratory investigations. Each blood sample was analyzed for CD4 T lymphocyte counts within 6 hours of sample collection. Each blood specimen was then spun at 500RPM for 5 minutes to enable separation of plasma. Thereafter, each plasma sample was tested for hepatitis B surface antigen (HBsAg), hepatitis B core IgM (HBcIgM), hepatitis C virus (HCV) and hepatitis E virus (HEV) by ELISA technique.

### Inclusion and Exclusion Criteria

Only healthy non-pregnant subjects who agreed to participatein the study were enrolled. Post enrolment, only participants without HBV, HCV or HEV infection were considered eligible for the CD4 T lymphocyte evaluation study. All participants with evidence of HBV, HCV or HEV infection were excluded from the study.

## Laboratory Analysis

### ELISA Screenings

#### HBsAg

A sandwich enzyme linked immunosorbent assay (ELISA) for detection of HBsAg (Diagnostic Automation/Cortez Diagnostic, California, USA) was used to screen all the 437plasma samples for the presence of HBsAg. The assay was carried out in accordance with manufacturer’s instructions.

#### HBcIgM

All the 437 plasma samples were screened using a two-step incubation, solid phase antibody capture ELISA for detection of anti-HBc IgM (Diagnostic Automation/Cortez Diagnostic, California, USA). The assay was carried out in accordance with manufacturer’s instructions.

#### HCV

Antibodies: A two-step incubation third generation indirectELISA technique for detection of HCV antibodies (Diagnostic Automation/Cortez Diagnostic, California, USA) was used to screen all the 437 plasma samples. The assay was carried out in accordance with manufacturer’s instructions.

#### HEV IgM

All the 437 plasma samples were screened using HEV IgM ELISA technique for the detection of HEV IgM antibody (Beijing Wantai Biological Pharmacy Enterprise Co. Ltd., China. The assay was carried out in accordance with manufacturer’s instructions.

For all the ELISA screens, the Emax endpoint ELISA microplate reader (Molecular Devices, California, USA) was used to determine the optical density after which the result was interpreted in accordance with the manufacturer’s instructions.

### Assay Procedures for the Cyflow® Counter flow cytometer

About 5mL of blood specimen was collected from each subject into appropriately labelled K2 EDTA, and properly mixed to avoid clot. Each blood specimen was mixed gently, and 20μL of whole blood was immediately dispensed into an appropriately labeledsample tube using graduated micropipette. Then, 20μL of CD4 monoclonal antibody was added, and content of the tube mixed gently for few seconds. The mixture was incubated at room temperature in the dark for 15minutes. Thereafter, 800μL of sample buffer was added and mixed properly. Then, sample was run using the Partec Cyflow (Sysmex Partec GmbH, Germany)and result recorded immediately.

### Data Collection and Analysis

Data were entered and analyzed using Statistical Package for Social Science (version 20.0). Data were analyzed using descriptive statistics, T-test and ANOVA at 0.05 level of significance.

### Ethical Considerations

Ethical clearance for this study was obtained from the Oyo State Ministry of Health (AD3/479/ 349). All procedures performed in the study werein accordance with the ethical standards of the institutional research committee and the Helsinki declaration.

## Results

Overall 37 (HBsAg=31; HBcIgM=7 (4+ 3participantspositive for both HBsAg and HBcIgM); HCV=2; HEV=0) participants who tested positive to either of the HBV markers, HCV or HEV antibodies were excludedin the study. A total of 400 (M=114; F=286; Mean age=34.45 ±16.3) participants who met the criteria (negative for HBsAg, HBcIgM, HCV and HEV) for enrolment into the study were included in the CD4 evaluation study (Table1).

**Table 1:** Summary of CD4 T Lymphocyte Count of the study population by Age and Gender.

| **Gender** | **Number (%)** | **Mean (SD)** | **Range** | **F-test** | **p-value** |
| --- | --- | --- | --- | --- | --- |
| Male | 114 (28.5) | 1046 (399) | 352-2510 | 0.667 | 0.415 |
| Female | 286 (71.5) | 1238 (398) | 328-2680 |  |  |
| **Total** | **400** | **1183 (407)** | **328-2680** |  |  |
| **Age Range (Year)** | **Number** | **Mean (SD)** | **Range** | **F-test** | **p-value** |
| <18 | 33 | 1073 (410) | 328-2510 | 3.083 | 0.010 |
| 18-25 | 144 | 1228 (399) | 352-2680 |  |  |
| 26-35 | 58 | 1300 (403) | 667-2494 |  |  |
| 36-45 | 56 | 1065 (327) | 530-1859 |  |  |
| 46-60 | 79 | 1175 (442) | 564-2545 |  |  |
| >60 | 30 | 1099 (426) | 560-2044 |  |  |
| **Total** | **400** | **1183 (407)** | **328-2680** |  |  |

### CD4 T Lymphocyte Cell Levels

Estimated mean CD4 T lymphocyte count of 1,183±407 (CD4 Range: 328-2680) cells/μl of blood was recorded for the study participants (Table 1). A total of 4 (1.0%), 151 (37.8%), 157 (39.2%), 74 (18.5), and 14 (3.5) of the study participants had CD4 T lymphocyte count range 352-500, 501-1,000, 1,001-1500, 1501-2,000, and >2,000 cells/μl of blood, respectively (Table 2). Further analysis of CD4 T cell count by gender showed an estimated mean CD4 count of 1,046±399 (Range: 352-2510) for the male, and slightly higher 1,238±398 (Range: 328-2680) for the femaleparticipants. Estimated mean CD4 T lymphocyte counts range from 1,073 (lowest) among the age group <18 years, and 1,300 (highest) among the age group 26-35 years (Table 1). Differences in the estimated mean CD4 count between different age groups varied significantly (P=0.010). Age range for the male was 9 to 87 years, while the female participants aged between 7 and 80 years. Analysis by occupation of participants showed that highest (46.3%) and lowest (0.5%) CD4 T lymphocyte levels were recorded among students and housewives, respectively (Table 3). No significant association was observed between CD4 T lymphocyte values and occupation.

**Table 2:** CD4 T Lymphocyte Profile of the study population by Gender.

| **CD4 COUNT** | **MALE (%)** | **FEMALE (%)** | **TOTAL (%)** |
| --- | --- | --- | --- |
| <500 | 1 (0.9) | 3 (1.0) | 4 (1.0) |
| 501-1000 | 63 (55.3) | 88 (30.8) | 151 (37.8) |
| 1001-1500 | 34(29.8) | 123 (43.0) | 157 (39.2) |
| 1501-2000 | 12 (10.5) | 62 (21.7) | 74 (18.5) |
| >2000 | 4 (3.5) | 10 (3.5) | 14 (3.5) |
| **TOTAL** | **114 (28.5)** | **286 (71.5)** | **400 (100)** |

**Table 3:** Range of CD4 T lymphocyte count of the study population by Occupation.

| **Occupation** | **Frequency (%)** | **CD4 T Lymphocyte Count** |
| --- | --- | --- |
| Civil Servant | 112 (28) | 555-2545 |
| Housewife | 2 (0.5) | 624-695 |
| Missionary | 4 (1.0) | 530-1228 |
| Retiree | 17 (4.3) | 561-2044 |
| Self-employed | 80 (20.0) | 560-2680 |
| Student | 185 (46.3) | 328-2510 |
| **Total** | **400 (100.0)** |  |

## Discussion

In Nigeria, CD4 T lymphocyte levels have been determined among populations in different regions, and attempt was made to establish NRVs. However, it is noteworthy that enrollees for the NRV study were defined as apparently healthy persons based only on their HIV negative status[18]. In this study, we estimated CD4 T lymphocyte cell values among healthy community dwellers who tested negative for HBV, HCV and HEV in Ibadan, and compared with the NRVs. We observed significantly higher CD4 T lymphocyte values among the study population in comparison to the NRVs.

Estimated mean CD4 T lymphocyte values in the study population were significantly higher (1,183 ±407 vs. 847 ±307) than the NRVs. Similarly, Adoga et al., [21]reported higher (1,030 ±367 vs. 847 ±307) estimated CD4 T lymphocyte values among participants free of other viral pathogensin addition to HIV. Thus, the observed higher CD4 T lymphocyte values recorded in these studies could be as a result of screening and exclusion of participants with other pathogens capable of interference. It could also be as a result of other factors previouslyreported to influence CD4 values. For example, factors including sex, age [11, 12], race [5, 10], diurnal rhythms [8, 22], physical and psychological stress, pregnancy, drug administration [9], tuberculosis, viral infections [16], presence of anti-lymphocyte auto antibodies and procedures such as splenectomy or strenuous physical activities [13, 23] have been shown to influence CD4 T lymphocyte counts in individuals. In addition, the role and characteristics of CD4 T lymphocyte cells in varied human viral infections including HIV [14] and hepatitis viruses [2, 3, 15-17, 19, 20] have been reported.

Higher estimated mean CD4 T lymphocyte cell counts were recorded in both male (1,046 ±399 vs. 782 ±272), and female (1,238 ±398 vs. 920 ±327) participants in this study than the

NRVs. Further, analysis of the results of this study by gender showed higher estimated mean CD4 T lymphocyte count for the female (1,238 ±398) in comparison to the male (1,046 ±399) participants. Though, difference in CD4 count was not statistically significant, it confirms previous results[5, 18, 21], and even among HIV-infected antiretroviral naïve subjects [24] that the females usually have higher CD4 T lymphocyte counts than the malecounterpart. However, it is pertinent to note that previous study by Zeh et al. [7] among rural community dwellers in Kenya observed no significant differences by gender.

Comparison of the estimated mean CD4 T lymphocyte counts of participants in this study by age shows statistically significant variation (p-value = 0.01). This is consistent with the NRVs [18], and in agreement with reports from previous studies [5, 21].Specifically, among different age groups we recorded higher CD4 T lymphocyte values than the NRVs in this study. This is also consistent with Adoga et al., [21], who reported higher than the NRVs. However, it differs from results of previous study by Zeh et al., [7] among rural community dwellers in Kenya who observed no significant differences by age.

## Conclusions

In this study, we estimated CD4 T lymphocyte cell values among healthy community dwellers in Ibadan who tested negative for HBV, HCV and HEV, and compared with the NRVs. We observed significantly higher CD4 T lymphocyte values among the study population in comparison to the NRVs. Thus, we advise careful interpretation and use of extrapolated CD4 T lymphocyte values in the management of persons with diverse geographical background or healthconditions. The authors are not unaware of the significance of HIV status in CD4 T lymphocyte counts, however, a major limitation of this study was our inability to screen the study participants for detectable HIV antigens or antibodies. Doing so will violate our agreement to screen them for hepatitis viruses alone.However, it is noteworthy that participants wereexamined for any clinical indications of early HIV infection, thus, level of bias in the results could be said to be negligible.

